# A programmable system to methylate and demethylate m^6^A on specific mRNAs

**DOI:** 10.1101/2021.04.16.440100

**Authors:** Chen Chang, Gang Ma, Edwin Cheung, Andrew P. Hutchins

## Abstract

RNA N^6^-Methyladenosine (m^6^A) is the most abundant mRNA modification, and forms part of an epitranscriptomic system that modulates RNA function. RNA modifications can be reversibly catalyzed by several specific enzymes, and those modifications can be recognized by RNA binding proteins that in turn regulate biological processes. Although there are many reports demonstrating m^6^A participation in critical biological functions, this exploration has mainly been conducted through the global knockout or knockdown of the writers, erasers, or readers of m^6^A. Consequently, there is a lack of information about the role of m^6^A on single transcripts in biological processes, posing a challenge in understanding the biological functions of m^6^A. Here, we demonstrate a CRISPR/dCas13a-based RNA m^6^A-editor which can target mRNAs using single crRNA or multiple crRNAs array to methylate or demethylate m^6^A. We systematically assay its capabilities to enable the targeted rewriting of m^6^A dynamics, including modulation of circular RNA translation and transcript half-life. Finally, we demonstrate the utility of the system by specifically modulating XIST m^6^A levels, which can control X chromosome silencing and activation. Based on our editors, m^6^A on single and multiple transcripts can be modified to allow the exploration of the role of m^6^A on in biological processes.

## Introduction

Analogous to the DNA epigenetic system, RNA is also chemically modified to impart epitranscriptomic information. N^6^-methyladenosine (m^6^A) is the most abundant RNA chemical modification in higher eukaryotes, and marks all classes of mRNA, including coding, and long and short noncoding transcripts. In a typical cell type in mammals, several thousand transcripts are extensively marked by m^6^A^1,2^, generally in the 3’UTR in coding transcripts^1^, but anywhere in noncoding transcripts^3^. Increasingly, m^6^A has been implicated in a diverse range of cellular processes, centered on RNA metabolism and processing, including RNA stability, alternative splicing, nuclear export, retrotransposon silencing, and protein translation^4–14^. Additionally, m^6^A has also been recognized to play a role in several biological phenotypes, including embryonic stem cell differentiation^9,15^; cancer^5,16,17^, neurobiology^18^, and X chromosome inactivation^19^, amongst many others^16,20–22^. However, these mechanisms have been largely explored through the global knockout or knockdown of m^6^A effectors, such as the writers (METTL3/METTL14/WTAP), erasers (FTO/ALKBH5) or the readers (YTH protein-family)^9,12,14,21,23^. Considering m^6^A marks several thousand transcripts in a typical cell type, the scope for pleiotropic effects from these knockdowns/knockouts is large. Hence, there is a need to specifically modulate m^6^A on single or several transcripts to reveal the independent functions of m^6^A.

The Cas9 system of bacterial proteins utilize single guide RNAs to specifically cleave invading DNA and RNA pathogens^24,25^. This bacterial immune system has been repurposed to create a suite of powerful genome and epigenome modification tools^26–28^. Whilst Cas9 targets DNA, bacteria also have a system that specifically targets RNA, Cas13a, that degrades RNA with high efficiency^29–31^. Similar to the nuclease mutated dCas9, that has been used to target specific sequences of DNA to great effect^26,27^, there is also a nuclease mutated dCas13a that can serve the same purpose, and deliver fusion proteins to specific RNAs^32^. Thus, catalytic dead dCas13a, fused to RNA modifying enzymes, and with specific crRNAs can potentially modulate the epitranscriptome on single transcripts. To date, m^6^A editing systems have been described for the writing of m^6^A, using dCas13b-METTL3^33^, and a second system based on a dCas9-METTL3-METTL14^34^, and a third system based on CasRx-ALKBH5^35^. Here, we generated dCas13a fused with full-length METTL3, or the core m^6^A methyltransferase domain (MTD) of METTL3, or with the m^6^A demethylase FTO. We demonstrate our CRISPR/dCas13a associated RNA m^6^A editing system can be specifically navigated by crRNAs to modulate m^6^A on individual coding or non-coding RNAs. To explore the capabilities of these systems, we demonstrate the efficacy of this system to modulate circular RNA translational efficiency and transcript half-life. We show that our system can process a crRNA array containing multiple crRNAs to target several transcripts simultaneously for m^6^A editing. Finally, we utilize this system to edit m^6^A modification levels on the long-noncoding RNA (lncRNA) *XIST* in female cells, resulting in reactivation of *XIST*-repressed genes. This work describes a powerful tool to edit m^6^A levels on individual RNAs and helps expand the toolkit available to explore the impact of m^6^A on single or several specific transcripts.

## Results

### Construction of dCas13a associated m^6^A-editing systems in mammalian cells

To construct an m^6^A-editing system we started by fusing the catalytic inactivated LwCas13a (dCas13a)^29,30^ protein with RNA m^6^A methyltransferases or demethylase (**Fig. 1a, Supplementary Fig. 1a**). m^6^A methylation is catalyzed by several protein complexes, which contain multiple protein subunits, e.g., METTL3, METTL14, WTAP, etc.^22,36–41^. However, amongst these proteins METTL3 is the main catalytic enzyme for methylation processes, therefore, we fused human METTL3 full length protein, or a truncated methylation domain (MTD) to dCas13a as the m^6^A writers.

**Figure 1.**
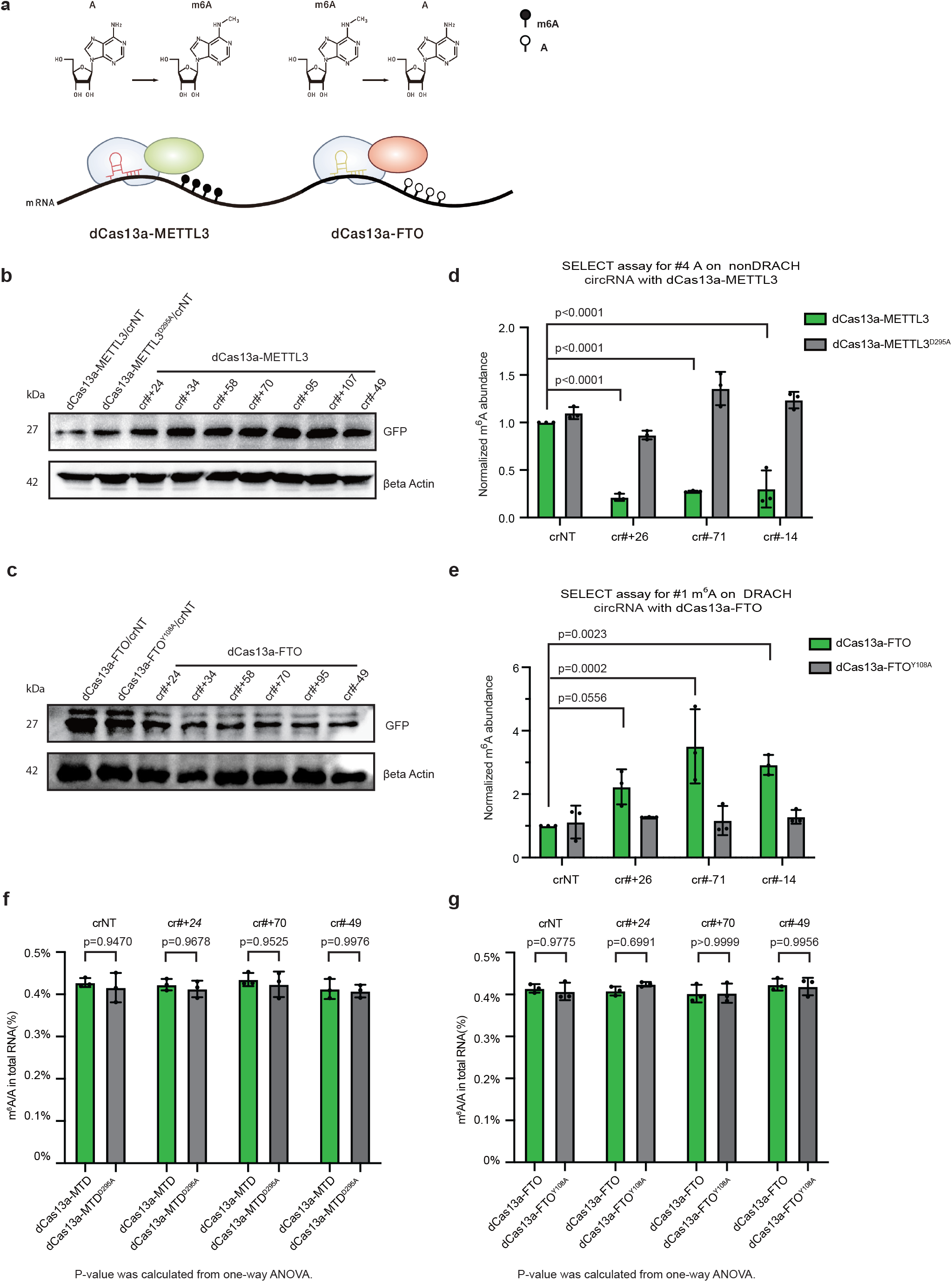
A programmable m^6^A methylation and demethylation system. (a) Schematic of the m^6^A editing systems (b) Western blot of GFP from 293T cells transfected with NT (non-targeting crRNA), or the crRNAs #1 thorough #6 targeting regions of the circular plasmid, and dCas13a-FTO or the catalytic dead dCas13a-FTO^D233A^. This experiment was repeated 3 times with similar results. (c) Western blot of GFP from 293T cells transfected with NT (non-targeting crRNA), or the crRNAs #1 thorough #6 targeting regions of the circular plasmid, and dCas13a-FTO, dCas13a-METTL3 or the catalytic dead dCas13a-METTL3^D395A^. This experiment was repeated 3 times with similar results. (d) SELECT assay of circGFPs m^6^A level, in cells transfected with dCas13a-METTL3, the catalytic null catalytic dead dCas13a-METTL3^D395A^ and crRNAs targeting the indicated base pairs upstream of the GFP ATG. Significance is from a two-tailed unpaired Student’s t-test, for this and all statistical tests used in the manuscript. Data is represented as the standard error of the mean (SEM), n=3 biological replicates with 3 technical replicates each. (e) SELECT assay of circGFPs m^6^A level, in cells transfected with dCas13a-FTO, the catalytic null catalytic dead dCas13a-FTO^Y108A^ and crRNAs targeting the indicated base pairs upstream of the GFP ATG. Data is represented as the SEM, n=3 biological replicates with 3 technical replicates each. (f) Whole transcriptome m^6^A/A ratio measurement for whole RNA m^6^A level in 293T cells transfected with the indicated crRNAs and dCas13a-METTL3, catalytic dead dCas13a-METTL3^D395A^. Data is represented as the SEM, n=3 biological replicates with 3 technical replicates each. (g) Whole transcriptome m^6^A/A ratio measurement for whole RNA m^6^A level in293T cells transfected with the indicated crRNAs and dCas13a-FTO or catalytic dead dCas13a-FTO^D233A^. Data is represented as the SEM, n=3 biological replicates with 3 technical replicates each.

Compared with m^6^A methylation, demethylation can be accomplished by a single catalytic enzyme^37,40–42^, such as FTO or ALKBH5, therefore, we fused full-length human FTO protein with dCas13a as an m^6^A eraser. As controls, we constructed catalytic-dead mutants of METTL3^D395A^ and FTO^D233A^ fused to dCas13a (**Supplementary Fig. 1a)**. All these fusion systems contain an N-terminal dCas13a, followed by a 13 amino acid GGS linker, a nuclear localization signal (NLS), to localize the fusion proteins in the nucleus, and a p2A with mCherry to report transfection efficiency. After transfection of the two systems into 293T cells mCherry fluorescence could be detected (**Supplementary Fig. 1b**), indicating expression in mammalian cells. The crRNA is contained on a second U6-promoter containing vector, which is co-transfected along with the fusion protein (**Supplementary Fig. 1a)**. Together these components make up an m^6^A editing system that can be targeted by the crRNA to a specific transcript.

### Fusion editing system can catalyze m^6^A residue changes on specific RNAs

To explore the activity of the dCas13a-fusions we initially constructed a reporter system to both directly and indirectly measure the activity m^6^A status of a targeted RNA. To this end, we exploited the ability of m^6^A to promote protein translation efficiency from circular RNAs in human cells^43–45^. To determine if our system can modify specific m^6^A sites on RNA, we modified a programmable circular RNA expression system to serve as a readout of m^6^A (**Supplementary Fig. 1c**)^45^. To this end, we generated an artificial circular RNA expression system where the coding sequence of GFP was separated linearly in two-parts between an IRES (Internal ribosome entry site), to form an ‘FP-IRES-G’ linear sequence that would circularize to form a complete GFP coding sequence (**Supplementary Fig. 1c**). Based on several reports^1,2,46^, in the endogenous setting, the METTL3/METTL14/WTAP complex prefers to methylate the A in an DRACH consensus motif, where D donates A, G or U, R donates A or G, and H donates A, C or U base pairs (**Supplementary Fig. 1d**). Hence, we inserted a sequence between the IRES and the ATG codon of GFP which includes ‘GGACU’ (DRACH) sequences that can be m^6^A methylated and promote circular GFP translation (**Supplementary Fig. 1c, e**). Circular RNAs containing 2, 1 or no GGACU sequences had decreasing levels of GFP translation, as measured by Western blot (**Supplementary Fig. 1e**). In the endogenous METTL3/METTL14 complex, METTL14 targets METTL3 to DRACH motifs^1,2,46^, however, as our system does not use METTL14, the requirement for DRACH targeting is relaxed, and potentially any available A base pair is a target. Consequently, we used the No RRACH containing FP-G circular reporter, with several extra A base pairs inserted into the region between the terminal of IRES sequence, to act as potential m^6^A sites (**Supplementary Fig. 1f**). We then selected several sites just outside the IRES region, before or after the start codon of the GFP, and designed crRNAs containing the complementary sequences to those selected sites (**Supplementary Fig. 1f**). We then co-transfected the crRNA with dCas13a-METTL3 or dCas13a-FTO, and performed Western blot assay for GFP. Western blot showed that, compared with non-targeting (NT) crRNA, crRNAs targeting the circular RNA resulted in improved GFP translation for the dCas13a-METTL3 constructs, and impaired translation for the dCas13a-FTO system (**Fig. 1b, c and Supplementary Fig. 1g, h**). This effect was lost when the catalytic null METTL3^D295A^ and FTO^Y108A^ were used (**Fig. 1b, c and Supplementary Fig. 1g, h**). Interestingly, little bias was introduced if the crRNA was more, or less, distal from the IRES and ATG of GFP, and even up to 107 bp 5’ of the ATG it was capable of promoting GFP translation, in the case of dCas13a-METTL3, or up to 93 bp 5’ of the ATG for dCas13a-FTO (**Fig. 1b, c**) We then performed an assay based on competitive elongation and ligation-based qPCR to measure the level of m^6^A at a specific residue (SELECT assay)^47^. In the SELECT assay, the presence of m^6^A impairs the production of the cDNA product, and increased m^6^A manifests as a decreased cycle threshold, and decreased m^6^A the inverse^47^. The SELECT assay showed that dCas13a-METTL3 with specific crRNAs resulted in an up-regulation in the level of m^6^A (**Fig. 1d)** and the up-regulation was lost if the catalytic mutant forms of METTL3^D295A^ was used (**Fig. 1d**). Conversely, dCas13a-FTO resulted in an increase in m^6^A, which was also lost in the FTO^Y108A^ catalytic mutant (**Fig. 1e**). One concern is that the overexpression of dCas13a-METTL3/FTO may act as a dominant positive, and non-specifically methylate RNA. ELISA for the transcriptome-wide level of m^6^A/A ratio indicated that the dCas13a fusions did not significantly change the transcriptome level of m^6^A, using either the active or catalytic null versions of METTL3/FTO (**Fig. 1f, g**).

### dCas13a-METTL3, dCas13a-MTD and dCas13a-FTO fusions can modulate m^6^A on specific endogenous transcripts, and alter their half-life

We next set out to test the fusion proteins on endogenous transcripts. To verify whether m^6^A editors can edit m^6^A on endogenous transcripts, we choose several candidates, including mRNAs (*H1F0, SGK1*) and lncRNAs (*MALAT1, ID3*). Reanalysis of m^6^A abundance data in 293T cells indicated these transcripts are expressed and marked by m^6^A in the normal state (**Supplementary Fig. 2a-d**)^1,48^. Using a crRNA targeting *SGK1*, SELECT assay indicated that m^6^A levels were significantly increased in dCas13a-MTD, or decreased in dCas13a-FTO on the *SGK1* RNA, but were unaltered with the non-targeting crRNA, or with the METTL3 or FTO catalytic nulls (**Fig. 2a, b**). dCas13a-MTD, cotransfected with a crRNA targeting *H1F0* showed a similar effect (**Fig. 2c**), and dCas13a-FTO with the same crRNA against *H1F0* led to a significant decrease in m^6^A levels (**Fig. 2d**). RIP (RNA immunoprecipitation)-qPCR using an antibody against m^6^A indicated dCas13a-MTD significantly increased m^6^A levels on *H1F0* transcripts (**Fig. 2e**), whilst dCas13a-FTO led to a significant (albeit modest) decrease in m^6^A on *H1F0* or *ID3* transcripts, when the respective crRNAs were co-transfected (**Fig. 2f**). We next explored the effect on the lncRNA *MALAT1*. As for coding transcripts, dCas13a-MTD significantly increased m^6^A levels, as measured by both SELECT, and m^6^A RIP-qPCR (**Supplementary Fig. 3a, b**), whilst dCas13a-FTO significantly reduced m^6^A (**Supplementary Fig. 3c, d**). Importantly, SELECT indicated the modulation of m^6^A was only achieved when using catalytically active dCas13a-METTL3/FTO (**Supplementary Fig. 2a, c**). Finally, to confirm that the dCas13a-MTD/FTO fusions are not acting as dominant positives, we measured the whole transcriptome level of m^6^A/A ratio, which indicated there was no significant changes (**Supplementary Fig. 2e, f**). These assays indicate that m^6^A editors can modify m^6^A on specific target transcripts.

**Figure 2.**
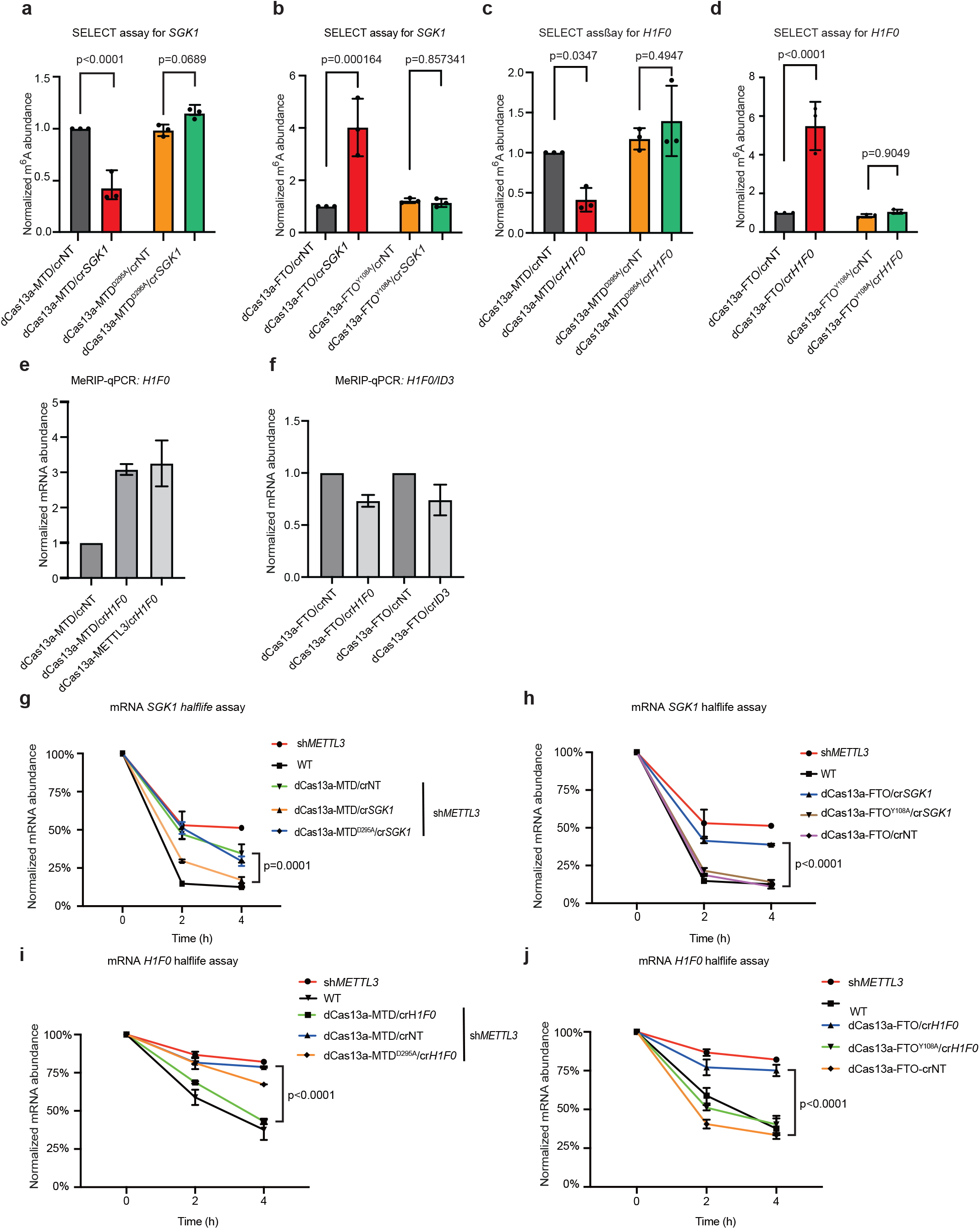
m^6^A editors can alter the half-life of specific endogenous transcripts. (a) SELECT assay for m^6^A level on *SGK1*, with dCas13a-MTD and catalytic null and crRNA targeting *SGK1*. Data is represented as the SEM, n=3 biological replicates with 3 technical replicates each. (b) SELECT assay for m^6^A level on *SGK1*, with dCas13a-FTO and catalytic null and crRNA targeting *SGK1*. Data is represented as the SEM, n=3 biological replicates with 3 technical replicates each. (c) SELECT assay for m^6^A level on *H1FO*, with dCas13a-MTD and catalytic null and crRNA targeting *H1FO*. Data is represented as the SEM, n=3 biological replicates with 3 technical replicates each. (d) SELECT assay for m^6^A level on *H1FO*, with dCas13a-FTO and catalytic null and crRNA targeting *H1FO*. Data is represented as the SEM, n=3 biological replicates with 3 technical replicates each. (e) MeRIP-qPCR assay, with dCas13a-MTD co-transfected with crRNA against *H1FO*. Data is represented as the SEM, n=3 biological replicates with 3 technical replicates each. (f) MeRIP-qPCR assay, with dCas13a-FTO co-transfected with crRNAs against *H1FO or ID3*. Data is represented as the SEM, n=3 biological replicates with 3 technical replicates each. (g) *SGK1* half-life assay, with dCas13a-MTD co-transfected with crRNA against *SGK1*. Data is represented as the SEM, n=3 biological replicates with 3 technical replicates each. (h) *SGK1* half-life assay. With dCas13a-FTO co-transfected with crRNA against *SGK1*. Data is represented as the SEM, n=3 biological replicates with 3 technical replicates each. (i) *H1F0* half-life assay, with dCas13a-MTD co-transfected with crRNA against *H1F0*. Data is represented as the SEM, n=3 biological replicates with 3 technical replicates each. (j) *H1F0* half-life assay. With dCas13a-FTO co-transfected with crRNA against *H1F0*. Data is represented as the SEM, n=3 biological replicates with 3 technical replicates each.

**Figure 3.**
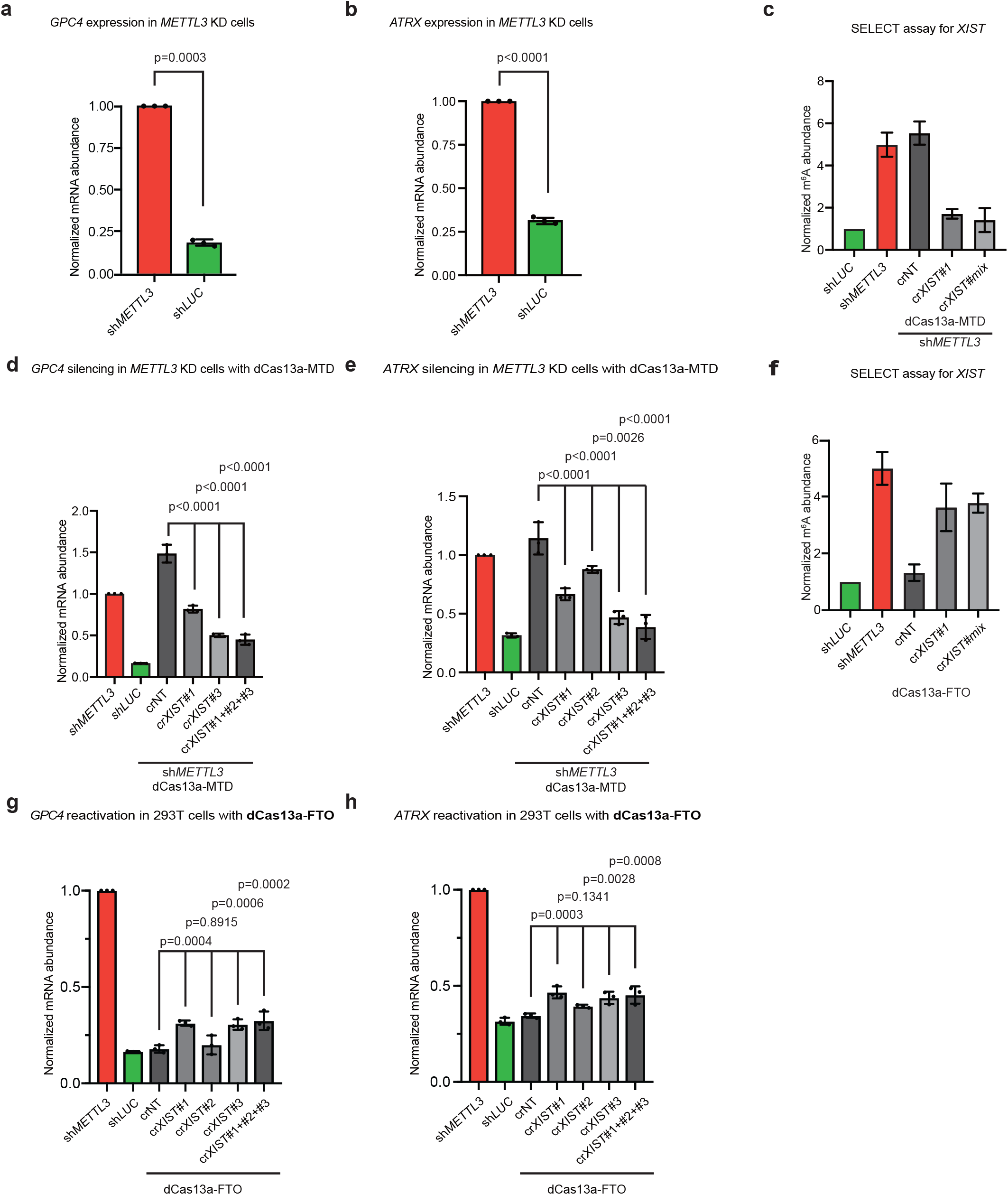
m^6^A editors can modulate *XIST* function and reactivate X chromosome inactivated genes. (a) Expression of the X chromosome expressed gene *GPC4* when *METTL3* was knocked down in 293T cells. Data is represented as the SEM, n=3 biological replicates with 3 technical replicates each. (b) Expression of the X chromosome expressed gene *ATRX* when *METTL3* was knocked down in 293T cells. Data is represented as the SEM, n=3 biological replicates with 3 technical replicates each. (c) SELECT assay for m6A levels on *XIST*, when METTL3 was knocked down, compared to cells with dCas13a-MTD and crRNAs targeting one or several regions of *XIST*. Data is represented as the SEM, n=3 biological replicates with 3 technical replicates each. (d) Expression of *GPC4* when *METTL3* was knocked down, compared to cells co-transfected with dCas13a-MTD and crRNAs targeting one or several regions of *XIST*. Data is represented as the SEM, n=3 biological replicates with 3 technical replicates each. (e) Expression of *ATRX* when *METTL3* was knocked down, compared to cells co-transfected with dCas13a-MTD and crRNAs targeting one or several regions of *XIST*. Data is represented as the SEM, n=3 biological replicates with 3 technical replicates each. (f) SELECT assay for m^6^A levels on *XIST*, when METTL3 was knocked down, compared to sh*LUC* transfected cells with dCas13a-FTO and crRNAs targeting one or several regions of *XIST*. Data is represented as the SEM, n=3 biological replicates with 3 technical replicates each. (g) Expression of *GPC4* when *METTL3* was knocked down, compared to cells co-transfected with dCas13a-FTO and crRNAs targeting one or several regions of *XIST*. Data is represented as the SEM, n=3 biological replicates with 3 technical replicates each. (h) Expression of *ATRX* when *METTL3* was knocked down, compared to cells co-transfected with dCas13a-FTO and crRNAs targeting one or several regions of *XIST*. Data is represented as the SEM, n=3 biological replicates with 3 technical replicates each.

RNA half-life increases when m^6^A writers or readers are knocked-down or knocked-out^9,49^. However, as these results have been demonstrated mainly in global knockout or knockdowns of m^6^A effectors, we wondered if our system could be used to alter the half-life of specific transcripts by changing the level of m^6^A. In our experiments, we used shRNA to knockdown *METTL3* in 293T cells, and compared the knockdowns to cells transfected with dCas13a-MTD or dCas13a-FTO, and then measured RNA half-life by treating cells with Actinomycin D, and then measuring RNA abundance by qRT-PCR. We chose two RNAs, *SGK1* and *H1F0*, as their transcript half-life has been previously characterized as being controlled by m^6^A^49^. The results showed that dCas13a-MTD, in cells with *METTL3* knockdown, can significantly increase the stability of the targeted RNAs, and reverted their stability close to that of the wildtype levels of *SGK1* and *H1F0* (**Fig. 2g, h**). dCas13a-FTO, with crRNAs against *SGK1* or *H1F0* significantly decreased the respective mRNA half-lives, and brough them close to the half-life in the METTL3 knockdown cells (**Fig. 2i, j)**. In all cases the catalytic null of MTD or FTO could not mediate these effects (**Fig. 2g-j**). These results shows that dCas13a-MTD and dCas13a-FTO can modulate m^6^A levels and impact transcript half-life.

### X chromosome-suppressed genes can be reactivated by modulating m^6^A level on XIST

We next set out to utilize our methylation modulating systems to alter a biological process. Only one X chromosome in female somatic cells is active, and the inactivated X chromosome is silenced through the activity of the non-coding RNA *XIST* (X-inactive specific transcript), which is expressed during development and in somatic cells and silences one X chromosome^50–53^. *XIST* is expressed and then coats one of the two X chromosomes, recruits polycomb and other epigenetic suppressors and methylates histones to silence one X chromosome in a complex regulatory process^54^. The *XIST* lncRNA itself is heavily m^6^A methylated, and when m^6^A readers were knocked down, X chromosome genes were reactivated^4^. These results suggest that m^6^A plays a critical role in X chromosome inactivation, however as m^6^A marks many thousands of transcripts, the precise role of m^6^A on *XIST*, and whether it is a determinant for X chromosome silencing is unclear. 293T cells have multiple X chromosomes^55^, however, all are silenced except one active X^54^, and consequently they can serve as a model for X chromosome silencing^54^. When we knocked down *METTL3* in 293T cells the X chromosome genes *GPC4* and *ATRX* were reactivated (**Fig. 3a, b**), in agreement with a previous report^4^. We designed several *XIST* targeting crRNAs against the three major sites of m^6^A in the *XIST* transcript (**Supplementary Fig. 4a**). We then co-transfected crRNAs targeting *XIST*, with dCas13a-MTD in *METTL3* knockdown 293T cells (**Supplementary Fig. 4b**). SELECT assay confirmed that the dCas13a-METTL3 system had methylated the *XIST* mRNA (**Fig. 3c**), and qPCR results showed that *ATRX* and *GPC4* were significantly repressed by dCas13a-MTD/cr*XIST*#1 and dCas13a-MTD/cr*XIST*#3 (**Fig. 3d, e**). However, cr*XIST*#3 seemed to be more effective, and a co-transfection of a mixture of all three crRNAs targeting *XIST* produced a modest improvement (**Fig. 3d, e**). Finally, transfection of the dCas13a-FTO/cr*XIST*s into 293T cells led to demethylation of m^6^A on *XIST*, as confirmed by SELECT assay (**Fig. 3f)** and a modest (but significant) reactivation of *GPC4* and *ATRX* (**Fig. 3g, h**). Presumably these effects are not so potent as demethylating *XIST* alone can only partially overcome the polycomb induced block. These results indicate that dCas13a-MTD and dCas13a-FTO can specifically alter the level of m^6^A on *XIST*, and lead to a change in the X chromosome activation state.

**Figure 4.**
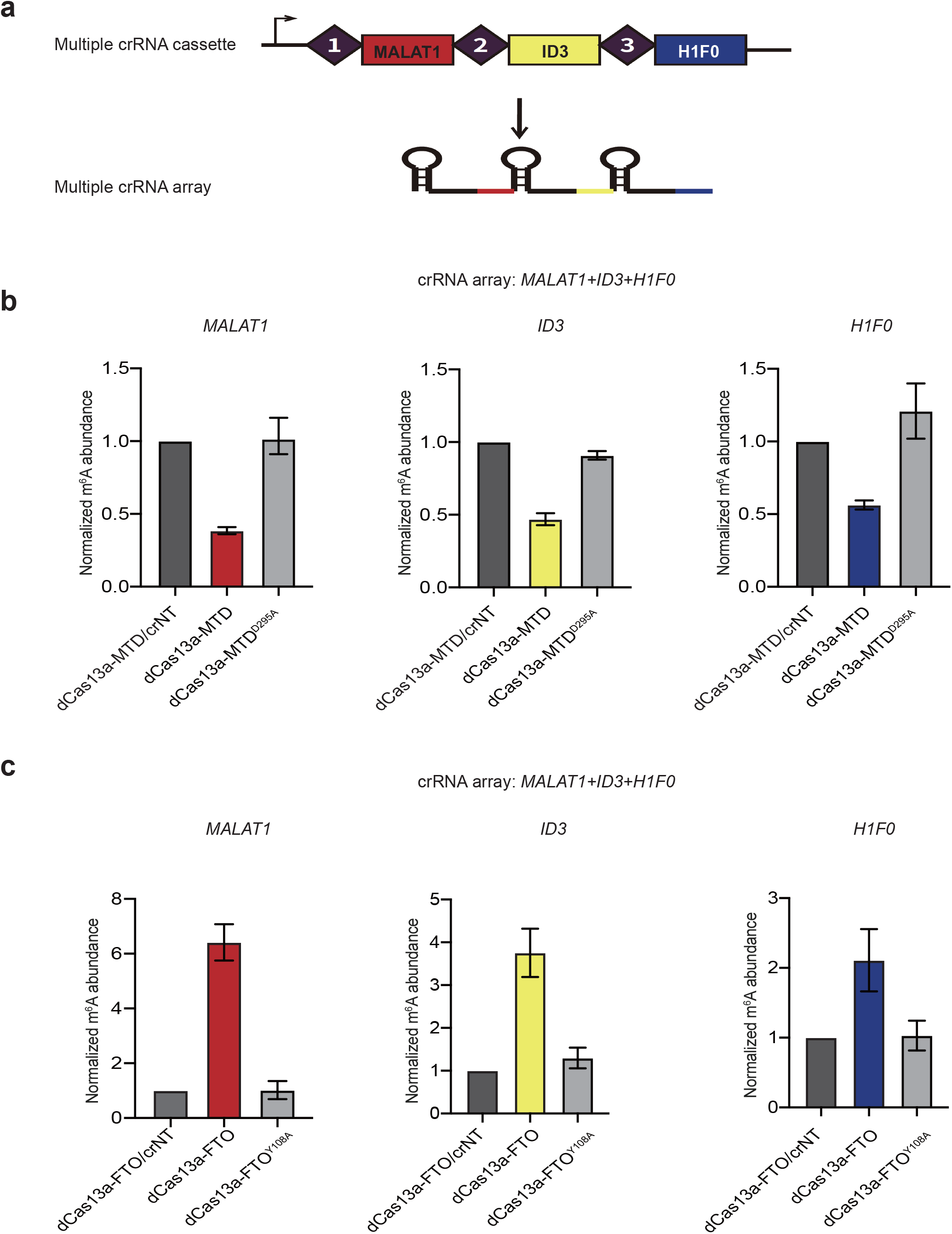
dCas13a can process a multiple crRNA cassette for targeting multiple transcripts. (a) Schematic of the multiple crRNA cassette with crRNAs targeting *MALAT1, ID3* and *H1F0*. The mature array will produce 3 crRNAs. (b) SELECT assay for the normalized m^6^A abundance of the three genes in the crRNA array, co-transfected with dCas13a-MTD or the catalytic null dCas13a-MTD^D295A^, with the non-targeting crNT as a control, or with the crArray from **panel a**. m^6^A level is normalized to the crNT transfection. Data is represented as the SEM, n=3 biological replicates with 3 technical replicates each. (c) As in **panel b**, but using dCas13a-FTO and the catalytic null dCas13a-FTO^Y108A^.

### System for multiple editing of m^6^A mRNAs

As we have shown with the above *XIST* experiments, many mRNAs have multiple hotspots for m^6^A, and potentially several sites must be simultaneously targeted to achieve a biological effect (**Supplementary Fig. 4a, Fig. 3d, e, g, h**). Additionally, genes can have multiple splice variants, and different genes’ transcripts can work in groups, hence it would be beneficial to modulate several transcripts m^6^A levels simultaneously. In Cas13a, the RNase domain and the crRNA maturation domains are located on different sites of the full length of Cas13a, in our study, we mutated the RNase domain on Cas13a, but not crRNA maturation domain, hence we investigated if dCas13a can mature a pre-crRNA array to functional crRNAs. We constructed a crRNA array containing multiple crRNAs, driven by a U6 promoter driving expression of the crRNA array, which can then be processed by dCas13a to produce separate crRNAs (**Fig. 4a**). Expression of this vector in 293T cells with crRNAs against *MALAT1, ID3* and *H1F0*, all led to a significant increase in m^6^A level on all 3 transcripts, as measured by SELECT assay, and the effect was lost if the catalytic null dCas13a-MTD was used (**Fig. 4b**). dCas13a-FTO was also capable of processing the crRNA array and could simultaneously significantly decrease the level of m^6^A in all three transcripts (**Fig. 4c**). Overall, we have described a system for the specific modification of m^6^A on multiple transcripts in an endogenous setting.

## Discussion

In this study, we have designed a CRISPR/dCas13a associated system to edit the m^6^A methylation of mRNA and lncRNAs. The catalytic dead Cas13a (dCas13a) was fused to either the full-length m^6^A methyltransferase protein METTL3 (dCas13a-METTL3), the catalytical domain of METTL3 (dCas13a-MTD), or the demethylase protein FTO (dCas13a-FTO), to alter m^6^A modification status of targeted RNA sequences. We first validated the systems by exploring the impact of dCas13a fusions on m^6^A levels and translation of GFP from an exogenous circular GFP reporter. Both dCas13a-METTL3/MTD and dCas13a-FTO could modulate m^6^A levels. We further show that these systems can also modulate m^6^A levels of linear endogenous transcripts, and these levels can impact transcript half-life. These results showed that our dCas13a based editors can alter the targeted RNA’s m^6^A modification level without disturbing the whole transcriptome m^6^A level^33^.

Several research groups have demonstrated a role of m^6^A on transcript half-life, however, a direct relationship between half-life and m^6^A has been challenging to show due to the widespread marking of transcripts with m^6^A and its role in many biological processes. Here, we utilized our m^6^A editors to altered several RNA’s m^6^A, half-life assay showed that targeted RNA’s half-life tends to be more stable when m^6^A level is decreased, and RNA tends to be less stable when m^6^A level is increased. As these experiments were performed in the absence of global knockouts or knockdowns they help establish a direct relationship between m^6^A levels and transcript half-life. Finally, we altered X-chromosome activation status by edited *XIST* RNA’s m^6^A. This is a critical result that demonstrates that m^6^A on *XIST* is crucial in enabling X chromosome suppression. Intriguingly, dCas13a-METTL3 could induce good resuppression of X chromosome genes, suggesting m^6^A on *XIST* is required for both initiation of X chromosome silencing. However, dCas13a-FTO led to a considerably more modest reactivation of X chromosome genes. Knockdowns of METTL3 can potently reactivate X chromosome genes (**Fig. 3a,b**)^19^, however our dCas13a-FTO/cr*XIST* experiment led to a significant, but modest, reactivation. Potentially, loss of m^6^A on *XIST* has only a weak ability to overcome other epigenetic suppression mechanisms. Alternatively, m^6^A must also be reduced on other transcripts to achieve robust X chromosome reactivation. Previous results used whole genome knockouts or knockdowns of m^6^A modulating enzymes, which will alter the m^6^A levels of several thousand transcripts which may indirectly modulate X chromosome inactivation. Here, we demonstrate that deposition of m^6^A on *XIST* alone is capable of silencing genes on the X chromosome, and removal of m^6^A on *XIST* can modestly reactive X chromosome genes.

It is increasingly clear there is an intimate interdependence between epigenetic control of DNA and epitranscriptomic control of RNA. For example, m^6^A marks the majority of DNA:DNA:RNA R-loops in stem cells^56^. m^6^A is emerging as a critical regulator of chromatin structure, transregulation of gene expression and stem cell differentiation^7,57,58^. The tool described here will be instrumental in dissecting these roles on specific individual transcripts and groups of transcripts. Our study has broad applications in the RNA targeting area, and based on our technique, the function of m^6^A on RNA can be precisely explored. We expect our dCas13a based editors can be applied to diverse cell types and biological settings to explore the action of m^6^A on specific transcripts. Utilizing this system researchers can isolate the effects of m^6^A on individual transcripts to explore pluripotency, development, cancer and human disease.

## Methods

### dCas13a-METTL3, MTD, FTO plasmid construction

dCas13a fused with METTL3, MTD, FTO separately, two NLS, one HA tag and p2A mCherry are added inside of each vector (**Supplementary Fig. 1a**).

### Cell culture and dCas13a-fusion transfection

Cells were maintained at 37°C with 5% CO2 in a humidified incubator and passaged every 2–3 days. Wild type HEK293T and HeLa cells were cultured in high-glucose Dulbecco’s Modified Eagle Medium (DMEM, ThermoFisher Scientific) supplemented with 10% FBS (Gibico). Cells were split with TrypLE Express (Life Technologies) according to the manufacturer’s instructions. At 80% confluency approximately 12h after plating, cells were transfected with 1,250 ng of dCas13a m6A editor plasmid and 1,250 ng of crRNA plasmid using 5 μl of Lipofectamine 3000 (Thermo Fisher Scientific) in Opti-MEM I Reduced Serum Media (Thermo Fisher Scientific). A full list of Cas13 crRNAs used in this work is given in **Supplementary Table 1**.

### RNA isolation and RT-qPCR

Total RNA was isolated from wild-type or transiently transfected cells with MiniBEST Universal RNA Extraction Kit (Takara), an additional DNase I (NEB) digestion step was performed to all samples to avoid DNA contamination, RNA concentration was measured by Nanodrop (Thermo Scientific). First-strand cDNA was synthesized by reverse transcription of 1 μg RNA using PrimeScrip RT Master Mix(Takara). Quantitative real time-PCR was performed using TB Green Premix Ex Taq (Takara) in QuantStudio 7 Flex Real-Time PCR System (Life Technologies, USA). β-actin and GAPDH were used as reference genes for input normalization. The mRNA expression was measured by quantitative PCR using the ΔΔCT method. Primers for quantitative PCR are listed in **Supplementary Table 1**.

### METTL3 knockdown via shRNA

The short hairpin RNA (shRNA) targeting METTL3 used in this study was previously described^6^. At 80% confluency approximately 12h after plating, cells were transfected with 1,250 ng of dCas13a m6A editor plasmid and 1,250 ng of crRNA plasmid using 5 μl of Lipofectamine 3000 (Thermo Fisher Scientific) in Opti-MEM I Reduced Serum Media (Thermo Fisher Scientific). 48 hours after the first transfection, a second transfection was performed. Cells were maintained at 70–80% confluency and collected 96 h after the first transfection. Knockdown was confirmed by RTqPCR (**Supplementary Fig. 4b**).

### RNA half-life assay

HEK293T cells and mESCs for lifetime assay were cultured in 12-well plates, cultured cells were transfected with CAREER include different crRNAs and control separately at 50% confluency. After 12 h, each whole of 12-well plates was re-seeded into three 12-well plates, and each whole of plate was controlled to afford the same number of cells. After 48 h, actinomycin D was added to a concentration of 5 uM at 6 h, 3 h and 0 h before extracting total RNA by MiniBEST Universal RNA Extraction Kit (Takara). The abundances of the interest genes were detected measured in each time point by real-time quantitative PCR (qPCR) using GAPDH as the reference gene.

### SELECT assay

SELECT assay was performed as previously described^47^. Briefly, Total RNA was mixed with 40 nM Up Primer, 40 nM Down Primer and 5 μM dTTP in 17μl 1×CutSmart buffer (50 mM KAc (acetic acid), 20 mM Tris-HAc, 10 mM MgAc2, 100 μg/ml bovine serum albumin, pH 7.9) at RT. The RNA and primers were annealed by incubating the mixture using a temperature gradient: 1 min each at 90°C, 80°C, 70°C, 60°C, 50°C, and then 40°C for 6 mins. Subsequently, a 3 μl of mixture containing 0.01 U Bst 2.0 DNA polymerase, 0.5 U SplintR ligase and 10 nmol ATP was added to a final volume of 20 μl. The final reaction mixture was incubated at 37°C for 20 min, denatured at 95°C for 20 mins and kept at 4°C, until being subjected to RT-qPCR as described above. SELECT primers are listed in **Supplementary Table 1**.

### Western blot

Whole-cell extracts were extracted by directly lysing the cells with 1×RIPA Buffer (Beyotime), with 1 mM PMSF (Beyotime) added immediately before use. Samples were boiled by adding 6×sodium dodecyl sulphate (SDS) sample buffer for 10 min at 100°C and resolved using SDS-polyacrylamide gel electrophoresis. The proteins were probed with the following antibodies: Monoclonal anti-GFP (1:2000, Thermo Scientific), anti-*β-actin* (1:2000, Thermo Scientific). Immuno-detection was performed using HRP-conjugated Affinipure Goat Anti-Mouse IgG(H+L) (1:5000, SA0 001-1, Proteintech) or HRP-conjugated Affinipure Goat Anti-Rabbit IgG(H+L) (1:5000, SA00001-2, Proteintech) and ECL prime substrate (Bio-Rad) according to the manufacturer’s instructions.

### Whole transcriptome m^6^A measurements

Global m^6^A/m in total RNA was quantified by the EpiQuik m^6^A/m RNA Methylation Quantification Kit (Epigentek Group, Farmingdale, NY) following the manufacturers’ specifications and using 100 ng as input (in duplicate or triplicate).

### m^6^A-RIP-qPCR

Total RNA was fragmented in solution of 50 mM Tris-HCl, pH 8.0, 50 mM MgCl2 and heated at 95 °C for 8 min. The m^6^A-modified and unmodified control RNAs (New England Biolabs) were divided into fragmented RNA, and a portion was saved as input RNA. The remaining fragmented RNA was subjected to m^6^A immunoprecipitation based on the previously described m^6^A-seq protocol^59^ with several modifications: 30 μl of protein G magnetic beads (Thermo Fisher Scientific) were washed twice by IP reaction buffer (150 mM NaCl, 10 mM Tris-HCl, pH 7.5, 0.1% NP-40 in nuclease-free H2O), resuspended in 500 μl of reaction buffer, and tumbled with 5 μg of anti-m^6^A antibody (New England Biolabs) at 4°C overnight. After two washes in reaction buffer, the antibody–bead mixture was resuspended in 500 μl of the reaction mixture containing 10 μg of fragmented total RNA,100 μl of reaction buffer and 5 μl of RNasin Plus RNase Inhibitor (Promega), and incubated for at least 4 h at 4 °C. To remove unbound RNA, samples were washed 5× with each of the following buffers: reaction buffer (150 mM NaCl, 10 mM Tris-HCl, pH 7.5, 0.1% NP-40 in nuclease-free H2O), low-salt reaction buffer(50 mM NaCl, 10 mM Tris-HCl, pH 7.5, 0.1% NP-40 in nuclease-free H2O) and high-salt reaction buffer (500 mM NaCl, 10 mM Tris-HCl, pH 7.5, 0.1% NP-40in nuclease-free H2O). RNA was eluted in RLT buffer (Qiagen) and purified with RNA Clean & Concentrator-5 kits (Zymo Research). Purified RNA was reverse transcribed with High-Capacity RNA-to-cDNA (Thermo Fisher Scientific) according to the manufacturer’s protocol. The resulting cDNA was pre-amplified with SsoAdvanced PreAmp Supermix (Bio-Rad) using qPCR forward and reverse primers according to the manufacturer’s protocol. Then, qPCR was performed with IQ Multiplex Powermix (Bio-Rad). All reactions were performed and quantified on a CFX96 Real-Time PCR Detection System (Bio-Rad) in technical triplicate. For m^6^A target sites with multiple probes, the mean was taken for the resulting Ct qPCR values. To account for variability in RNA amounts, target Ct values were normalized to the geometric mean of the following internal controls: a methylated EEF1A1 control transcript, m^6^A-modified RNA spike-in controls (New England Biolabs) and a nontargeted region on the crRNA-targeted transcript. RIP-qPCR primers are listed in **Supplementary Table 1**.

### Statistical analysis and reproducibility

Significance is from an unpaired Student’s t-test, for all statistical tests used in the manuscript. Bar charts with error bars represent the standard error of the mean from at least 3 or more replicates. Significance was only calculated if the number of biological replicates was 3 or more.

## Acknowledgements

This work was supported by the National Natural Science Foundation of China (31970589), the Science and Technology Planning Project of Guangdong Province (2019A050510004), the Shenzhen Innovation Committee of Science and Technology grants (ZDSYS20200811144002008), the University of Macau (MYRG2020-00100-FHS) and the Macau Science and Technology Development Fund (0011/2019/AKP). Supported by the Center for Computational Science and Engineering of Southern University of Science and Technology.

