## Supplementary Figures for "A programmable system to methylate and demethylate m^6^A on specific mRNAs"

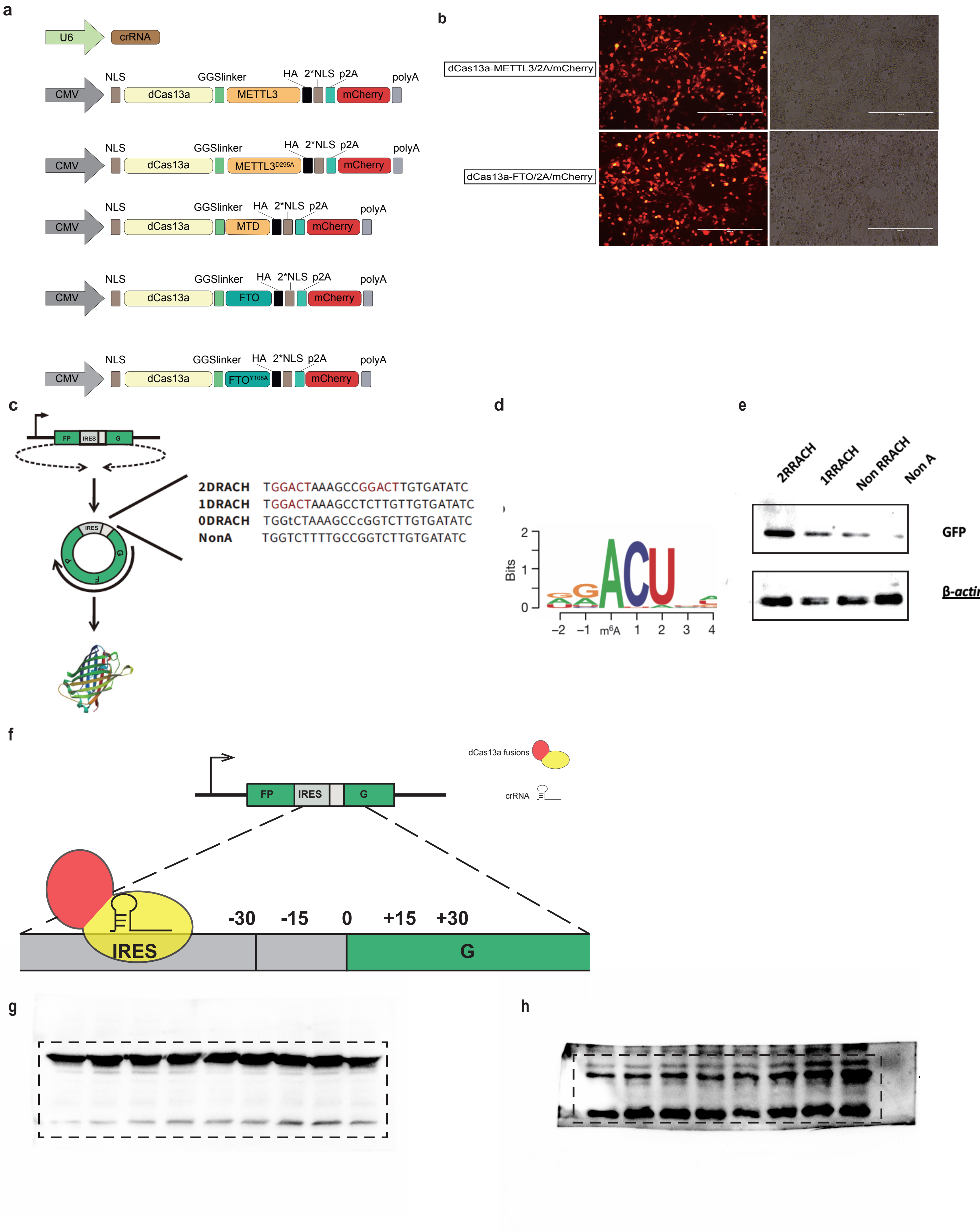


Supplementary Figure 1

**Supplementary Fig. 1. $m^6A$ editor systems can modulate the translation of circular RNAs.**

(a) Schematic of the constructs made in this manuscript.

(b) mCherry transfection images of the dCas13a-METTL3 or dCas13a-FTO constructs in 293T cells.

(c) Schematic of the circRNA construct.

(d) DRACH $m^6A$ position weight matrix motif.

(e) Western blot of GFP translated from circular RNA constructs containing 2, 1 or no DRACH motifs. This experiment was performed once.

(f) Schematic of the zoomed in region between the IRES and the ATG of GFP. Locations of the crRNAs are indicated by numbers, relative to the ATG (at base pair 0).

(g) Original, uncropped Western blot images for the panels in **Figure 1b, c**.

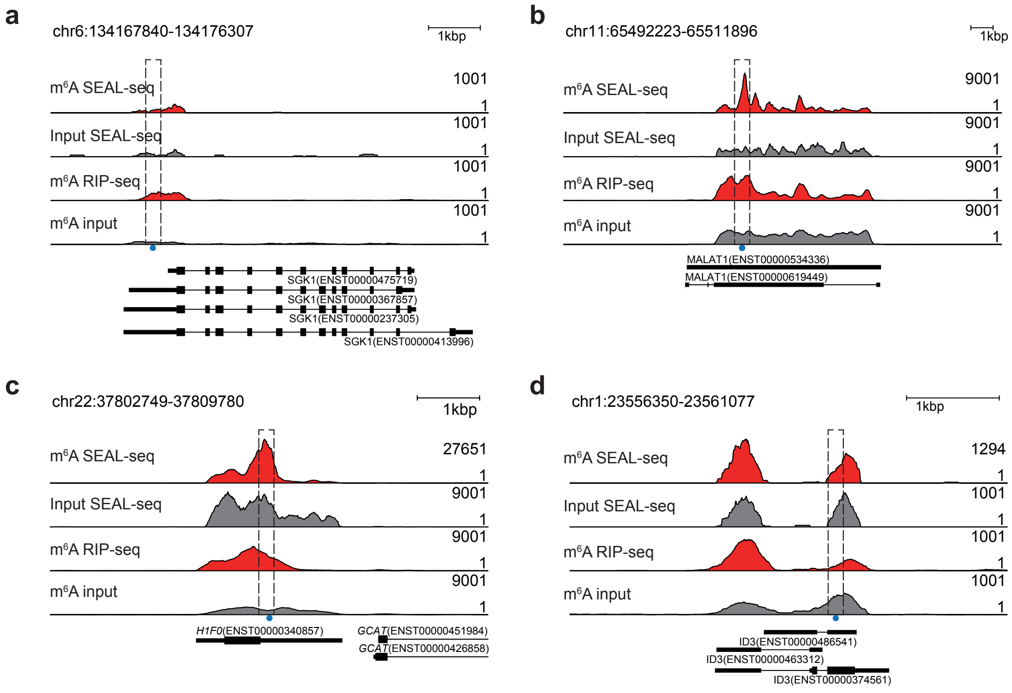


Supplementary Figure 2

**Supplementary Figure 2. Genome views of $m^6A$ levels of transcripts targeted by crRNAs in this study.**

(a) Genome view (hg38) of $m^6A$ RIP-seq data in 293T cells at the *SGK1* locus. The location of the crRNA is indicated with a grey bar. Red indicates $m^6A$ enrichment data, grey tracks indicate the corresponding input data. Transcripts are from GENCODE v32. $m^6A$ abundance data is from GSE129979[60] (top two rows) or GSE29714[1] (bottom two rows), for this and subsequent panels.

(b) As in panel a, but showing *MALAT1*

(c) As in panel a, but showing *H1F0*

(d) As in panel a, but showing *ID3*

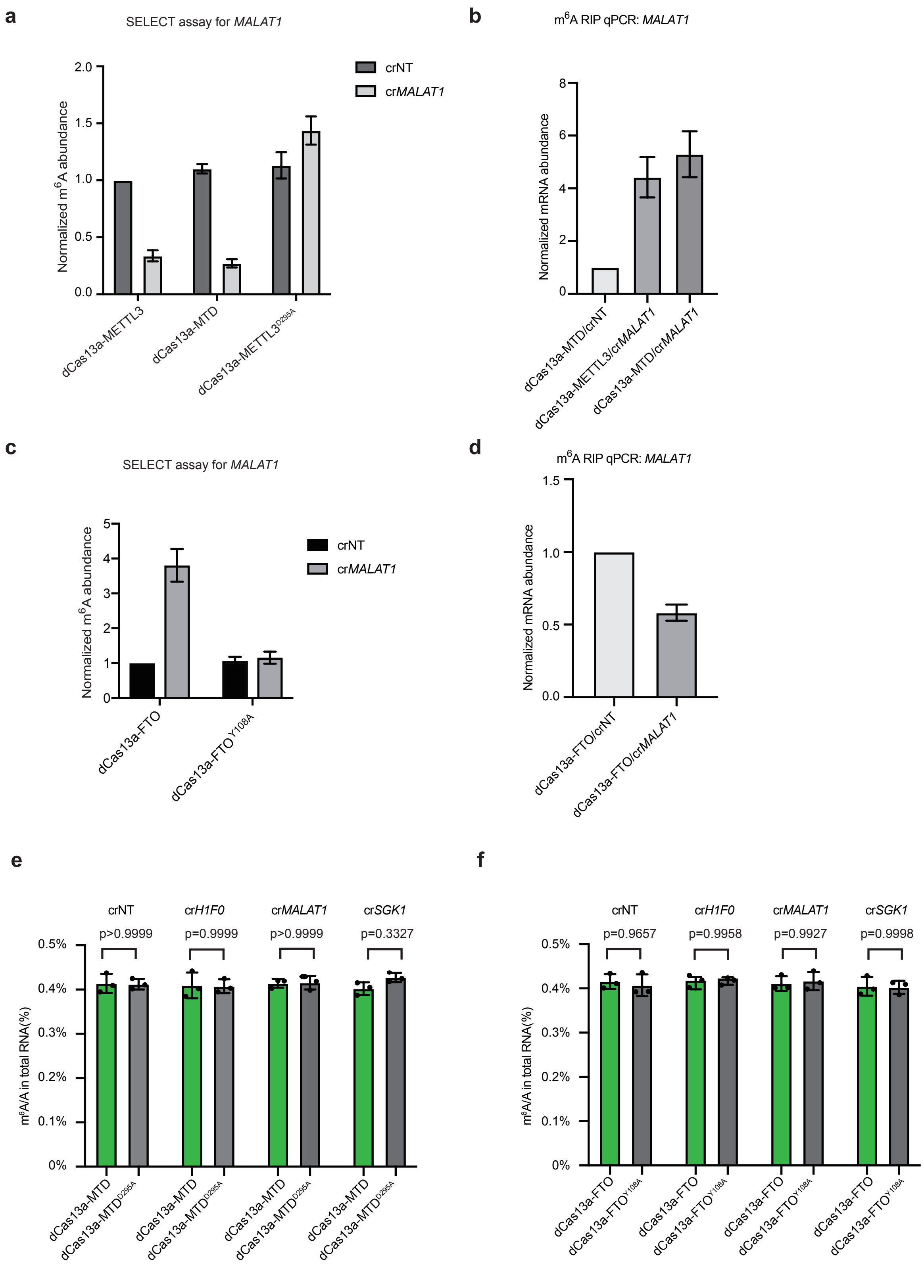


Supplementary Figure 3

**Supplementary Figure 3. m$^6$A editors can methylate and demethylate endogenous mRNA and long non-coding RNA transcripts.**

(a) SELECT assay for *MALAT1*, with dCas13a-MTD, dCas13a-METTL3 or the catalytic null form, with a non-targeting crRNA or with a crRNA targeting *MALAT1*. Data is represented as n=1 biological replicates with 3 technical replicates each.

(b) m$^6$A RIP-seq for *MALAT1* transcript, in cells transfected with dCas13a-METTL3, or dCas13a-MTD and non-targeting crRNA, or a crRNA targeting *MALAT1*. Data is represented as n=1 biological replicates with 3 technical replicates each.

(c) SELECT assay for *MALAT1*, with dCas13a-FTO or the catalytic null form, with a non-targeting crRNA or with a crRNA targeting *MALAT1*. Data is represented as n=1 biological replicates with 3 technical replicates each.

(d) m$^6$A RIP-seq for *MALAT1* transcript, in cells transfected with dCas13a- FTO, or dCas13a-MTD and non-targeting crRNA, or a crRNA targeting *MALAT1*. Data is represented as n=1 biological replicates with 3 technical replicates each.

(e) ELISA for ratio of m$^6$A/A in the indicated cells transfected with dCas13a-MTD, or its catalytic null, with the indicated crRNAs. Data is represented as the SEM, n=3 biological replicates with 3 technical replicates each.

(f) As in **panel e**, but using the dCas13a-FTO and its catalytic null.

**a**

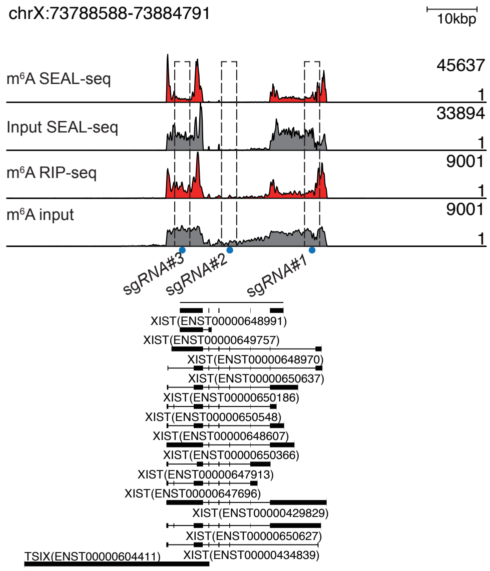


**b**

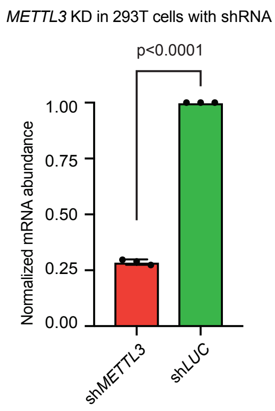


Supplementary Figure 4

**Supplementary Figure 4. $m^6A$ levels pattern on the *XIST* transcript and METTL3 knockdown.**

(a) Genome view of the $m^6A$ level in 293T cells on the *XIST* transcript. The location of the crRNAs are indicated with a grey bar. Transcripts are from GENCODE v32. $m^6A$ abundance data is from GSE129979[60] (top two rows) or GSE29714[1] (bottom two rows).

(b) Expression of *METTL3* after knockdown with shRNA. Data is represented as the SEM, n=3 biological replicates with 3 technical replicates each.
